## Supplementary Data for "ExhauFS: exhaustive search-based feature selection for classification and survival regression"

**Fig. S1. ROC curves for a toy cervical cancer dataset.** A: three best features found by ExhauFS ("perception_vulnerability", "socialSupport_instrumental", "empowerment_desires"). B: random forest classifier applied to the whole set of features. C: three most important features found by random forest classifier ("behavior_personalHygine", "empowerment_knowledge", "perception_severity"). Red points on ROC curves stand for the actual random forest classifier threshold values.

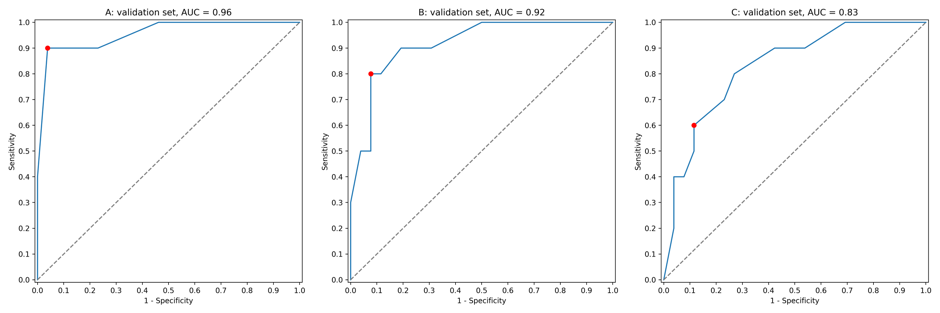

**Fig. S2. ROC and Kaplan-Meier plots for the four-gene signature (*TRIP13*, *ABAT*, *STC2*, *SIGLEC15*) evaluated on GSE1456 (microarray) and TCGA-BRCA (RNA-seq) datasets.** Red points on ROC curves stand for the actual SVM threshold values.

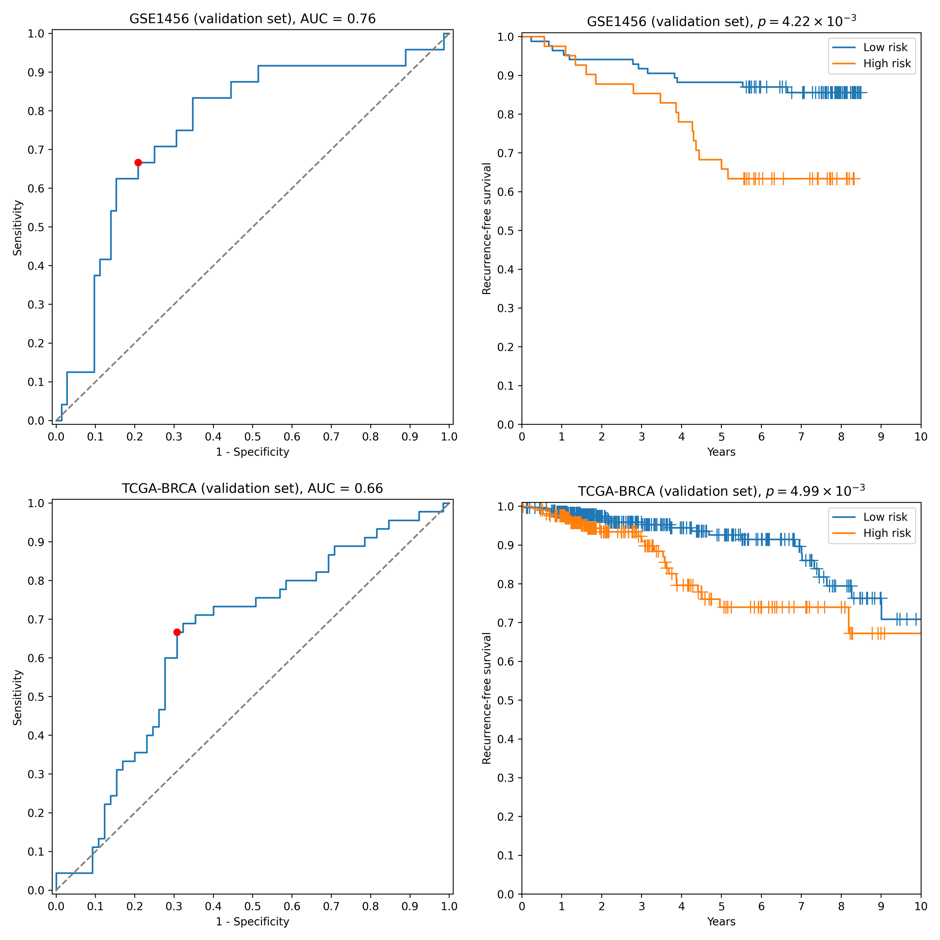

**Table S1. Numbers of patients in breast cancer datasets.**

| **Dataset** | **Number of patients with recurrence during the first 5 years** | **Number of recurrence-free patients with at least 7 years follow-up** | **Total number of patients** |
| --- | --- | --- | --- |
| GSE1456 | 24 | 72 | 126 |
| GSE3494 | 33 | 42 | - |
| GSE6532 | 30 | 49 | - |
| GSE12093 | 12 | 66 | - |
| GSE17705 | 26 | 120 | - |
| TCGA-BRCA | 45 | 66 | 559 |

**Table S2. Summary table for breast cancer prognostic classifiers constructed without feature pre-selection.** *N* filtration stands for number of classifiers which passed the filtration step, *N* validation stands for number of classifiers which additionally passed the same accuracy thresholds on the validation set.

| ***n*** | ***k*** | ***N* filtration** | ***N* validation** | **Percentage (validation / filtration)** |
| --- | --- | --- | --- | --- |
| 7762 | 1 | 0 | 0 | 0% |
| 1000 | 2 | 3 | 1 | 33.3% |
| 130 | 3 | 65 | 35 | 53.8% |
| 50 | 4 | 10 | 3 | 30% |
| 32 | 5 | 17 | 5 | 29.4% |
| 24 | 6 | 0 | 0 | 0% |
| 21 | 7 | 0 | 0 | 0% |
| 19 | 8 | 0 | 0 | 0% |
| 18 | 9 | 0 | 0 | 0% |
| 18 | 10 | 0 | 0 | 0% |
| 19 | 11 | 0 | 0 | 0% |
| 19 | 12 | 0 | 0 | 0% |
| 19 | 13 | 0 | 0 | 0% |
| 20 | 14 | 0 | 0 | 0% |
| 20 | 15 | 0 | 0 | 0% |
| 21 | 16 | 0 | 0 | 0% |
| 22 | 17 | 0 | 0 | 0% |
| 23 | 18 | 0 | 0 | 0% |
| 23 | 19 | 0 | 0 | 0% |
| 24 | 20 | 0 | 0 | 0% |

The last column of the table (total number of patients) stands for datasets used in Kaplan-Meier plots construction (Figure 2, Figure S2).

**Table S3. Summary table for breast cancer prognostic classifiers constructed with “stable” genes pre-selection.**

| ***n*** | ***k*** | ***N* filtration** | ***N* validation** | **Percentage (validation / filtration)** |
| --- | --- | --- | --- | --- |
| 7762 | 1 | 0 | 0 | 0% |
| 1000 | 2 | 0 | 0 | 0% |
| 130 | 3 | 15 | 4 | 26.7% |
| 50 | 4 | 265 | 90 | 34.0% |
| 32 | 5 | 1706 | 534 | 31.3% |
| 24 | 6 | 836 | 532 | 63.6% |
| 21 | 7 | 1410 | 1049 | 74.4% |
| 19 | 8 | 1686 | 1433 | 85.0% |
| 18 | 9 | 1498 | 1381 | 92.2% |
| 18 | 10 | 1773 | 1696 | 95.7% |
| 19 | 11 | 3607 | 3515 | 97.4% |
| 19 | 12 | 2990 | 2935 | 98.2% |
| 19 | 13 | 1879 | 1860 | 99.0% |
| 20 | 14 | 2779 | 2750 | 98.9% |
| 20 | 15 | 1272 | 1261 | 99.1% |
| 21 | 16 | 1343 | 1335 | 99.4% |
| 22 | 17 | 1672 | 1663 | 99.5% |
| 23 | 18 | 3470 | 3401 | 98.0% |
| 23 | 19 | 1156 | 1144 | 99.0% |
| 24 | 20 | 549 | 539 | 98.2% |

*N* filtration stands for number of classifiers which passed the filtration step, *N* validation stands for number of classifiers which additionally passed the same accuracy thresholds on the validation set.

**Table S4. Number of gene occurrences in signatures which passed the filtration step**.

| **Gene** | **Count** | **Percentage**  **(count / 1773)** | ***p*-value** | **Adjusted *p*-value** |
| --- | --- | --- | --- | --- |
| *ABAT* | 1666 | 94.0% | 1.20E-290 | **2.15E-289** |
| *CCNL2* | 1356 | 76.5% | 1.42E-75 | **1.28E-74** |
| *TRIP13* | 1256 | 70.8% | 2.48E-40 | **1.49E-39** |
| *EPN3* | 1251 | 70.6% | 6.83E-39 | **3.07E-38** |
| *IGFBP6* | 1158 | 65.3% | 2.61E-17 | **9.41E-17** |
| *ZWINT* | 1071 | 60.4% | 1.65E-05 | **4.95E-05** |
| *CX3CR1* | 1025 | 57.8% | 0.0262901 | 0.06760311 |
| *MB* | 1015 | 57.2% | 0.07231083 | 0.16269938 |
| *SLC7A5* | 946 | 53.3% | 0.96698141 | 1 |
| *ECHDC2* | 929 | 52.4% | 0.99595152 | 1 |
| *UBE2C* | 893 | 50.4% | 0.99999352 | 1 |
| *KIF4A* | 850 | 48.0% | 1 | 1 |
| *NUMA1* | 836 | 47.1% | 1 | 1 |
| *CFAP69* | 822 | 46.4% | 1 | 1 |
| *MTFR1* | 817 | 46.1% | 1 | 1 |
| *RTN1* | 799 | 45.1% | 1 | 1 |
| *STARD13* | 646 | 36.4354202 | 1 | 1 |
| *GINS2* | 394 | 22.2222222 | 1 | 1 |

Data is shown for n = 18, k = 10. Total number of considered gene signatures: 1773. Binomial test (1773 trials with success probability 10/18) was used to calculate *p*-values, multiple testing correction was done using Benjamini-Hochberg procedure.

**Table S5. Summary table for colorectal cancer prognostic regressors (concordance index-based feature selection).**

| ***n*** | ***k*** | ***N* filtration** | ***N* validation** | **Percentage (validation / filtration)** |
| --- | --- | --- | --- | --- |
| 86 | 1 |  | 0 | 0% |
| 86 | 2 |  | 0 | 0% |
| 86 | 3 |  | 0 | 0% |
| 59 | 4 | 332 | 27 | 8.1% |
| 37 | 5 | 1699 | 306 | 18.0% |
| 28 | 6 | 2954 | 720 | 24.4% |
| 24 | 7 | 7493 | 2435 | 32.5% |
| 22 | 8 | 8979 | 3874 | 43.1% |
| 21 | 9 | 8488 | 4558 | 53.7% |
| 20 | 10 | 5040 | 3413 | 67.7% |
| 20 | 11 | 3905 | 2988 | 76.5% |
| 20 | 12 | 2358 | 1966 | 83.4% |
| 20 | 13 | 1042 | 923 | 88.6% |
| 21 | 14 | 1072 | 937 | 87.4% |
| 22 | 15 | 1467 | 1247 | 85.0% |
| 22 | 16 | 383 | 341 | 89.0% |
| 23 | 17 | 773 | 711 | 92.0% |
| 23 | 18 | 134 | 129 | 96.3% |
| 24 | 19 | 150 | 135 | 90.0% |
| 25 | 20 | 67 | 60 | 89.6% |

*N* filtration stands for number of Cox models which passed the filtration step, *N* validation stands for number of models which additionally passed the same accuracy thresholds on the validation set.

**Table S6. Summary table for colorectal cancer prognostic regressors (feature selection based on the search of differentially expressed isomiRs between normal tissues and tumors).**

| ***n*** | ***k*** | ***N* filtration** | ***N* validation** | **Percentage (validation / filtration)** |
| --- | --- | --- | --- | --- |
| 86 | 1 | 0 | 0 | 0% |
| 86 | 2 | 0 | 0 | 0% |
| 86 | 3 | 191 | 42 | 22.0% |
| 59 | 4 | 1504 | 534 | 35.5% |
| 37 | 5 | 5292 | 1605 | 30.3% |
| 28 | 6 | 4441 | 1 | 0.0% |
| 24 | 7 | 11 | 7 | 63.6% |
| 22 | 8 | 70 | 50 | 71.4% |
| 21 | 9 | 113 | 94 | 83.2% |
| 20 | 10 | 105 | 96 | 91.4% |
| 20 | 11 | 87 | 82 | 94.2% |
| 20 | 12 | 47 | 42 | 89.4% |
| 20 | 13 | 23 | 20 | 87.0% |
| 21 | 14 | 5 | 4 | 80.0% |
| 22 | 15 | 20 | 18 | 90.0% |
| 22 | 16 | 4 | 4 | 100.0% |
| 23 | 17 | 1 | 1 | 100.0% |
| 23 | 18 | 0 | 0 | 0% |
| 24 | 19 | 0 | 0 | 0% |
| 25 | 20 | 120 | 15 | 12.5% |

*N* filtration stands for number of Cox models which passed the filtration step, *N* validation stands for number of models which additionally passed the same accuracy thresholds on the validation set.
